# Single-cell analysis of non-alcoholic fatty livers identifies a role for the constitutive androstane receptor

**DOI:** 10.1101/2022.08.15.504026

**Authors:** Laetitia Coassolo, Tianyun Liu, Yunshin Jung, Nikki P. Taylor, Meng Zhao, Gregory W. Charville, Silas Boye Nissen, Hannele Yki-Jarvinen, Russ Altman, Katrin J. Svensson

## Abstract

Non-alcoholic fatty liver disease is a heterogeneous disease with unclear underlying molecular mechanisms. While several genetic risk factors have been identified, the cellular and molecular heterogeneity associated with the development of hepatic steatosis are still not fully understood. Here, we perform single-cell RNA sequencing of hepatocytes and hepatic nonparenchymal cells to map the lipid signatures in mice with non-alcoholic fatty liver disease (NAFLD). We uncover previously unidentified clusters of hepatocytes characterized by either high or low *srebp1* expression with unique molecular signatures of lipid synthesis. We find that NAFLD livers have elevated expression of the constitutive androstane receptor (CAR), a gene previously associated with lipid synthesis. Furthermore, nuclear expression of CAR positively correlates with steatohepatitis in humans. These findings demonstrate significant cellular differences in lipid signatures and identify a gene with a high likelihood of being linked to hepatic steatosis in humans.

## INTRODUCTION

Non-alcoholic fatty liver disease (NAFLD) and the more severe condition non-alcoholic steatohepatitis (NASH), are now affecting 25 % of the global adult population (Diehl and Day, 2017). NAFLD is a progressive metabolic disease characterized by hepatic lipid accumulation, inflammation, fibrosis, and insulin resistance. NAFLD has become the most common form of chronic liver disease in the US and has no pharmacological treatment (Sanyal, 2019). In recent years, our molecular understanding of the process of diet-induced hepatic steatosis has emerged (Elizabeth M. Brunt et al., 2015; Diehl and Day, 2017). Several genes have been identified as risk factors for elevated hepatic lipid accumulation, including a single nucleotide polymorphism in the PNPLA3 gene, generating a PNPLA3-I148M variant that increases the risk of developing fatty liver disease (Romeo *et al*., 2008; Huang *et al*., 2010; Luukkonen *et al*., 2019) and an HSD17B13 variant that is protective (Abul-Husn *et al*., 2018; Luukkonen *et al*., 2020). In addition to genetic factors, environmental factors such as nutrient availability are well-known contributors to hepatic steatosis and NAFLD. Fat consumption (Tavares De Almeida *et al*., 2002; Donnelly *et al*., 2005; Puri *et al*., 2009), excessive fructose intake (Abdelmalek et al., 2010; Stanhope et al., 2009; Thuy et al., 2008), hyperinsulinemia and obesity (Gaggini et al., 2013; Luyckx et al., 2000) positively correlates with NAFLD. However, there is an unmet need to understand the molecular characteristics and changes in cell-type composition and expression in NAFLD.

Interestingly, earlier studies have shown that lipid accumulation is not uniformly induced in all hepatocytes, with most cells having a relatively low accumulation of lipids and fewer cells with very high lipid storage (Herms *et al*., 2013). Elevated intra-hepatocyte lipid content in NAFLD is partly attributable to increased *de novo* lipogenesis (Softic *et al*., 2017), a process largely driven by the transcription factor sterol regulatory element-binding protein-1 (SREBP-1) isoforms *srebp1a* and *srebp1c* (Kim and Spiegelman, 1996; Maxwell *et al*., 2003; Moon *et al*., 2012). Yet, whether *srebp1* is homogenously expressed in hepatocytes, and whether other transcriptional regulators are associated with NAFLD pathogenesis remains underexplored.

Here, by using single-cell analyses, we demonstrate cellular differences in lipid signatures in livers of mice with fatty liver disease. We find that both hepatocyte lipid accumulation and *srebf1* expression are heterogeneous and identify Nr1i3, the constitutive androstane receptor (CAR), as a gene that is enriched in NAFLD livers from mice. While prior reports on CAR have been conflicting (Stern, Kurian and Wang, 2022), we show that CAR protein levels localized to the nucleus are highly elevated in human livers of patients with steatohepatitis. Our study reveals how NAFLD alters the transcriptomic landscape of hepatocytes during liver pathogenesis and identifies a highly associated NAFLD gene in mice and humans.

## RESULTS

### Hepatic steatosis is characterized by high hepatocyte heterogeneity

Previous single-cell analyses of livers have shown remarkable heterogeneity in the cell populations within the liver (MacParland *et al*., 2018; Xiong *et al*., 2019). However, while hepatocyte proteomic analyses have shown dramatic subcellular reorganization of proteins during the development of fatty liver disease (Krahmer *et al*., 2018), the cellular and metabolic heterogeneity of hepatocytes in NAFLD remains largely uncharacterized.

To dissect the transcriptional changes upon induction of hepatic steatosis, we first set out to determine the earliest time point of detectable physiological and biochemical changes in liver cell populations isolated from mice fed a high-fructose, high-fat (NAFLD, or amylin) diet (Boland *et al*., 2019; Jiang *et al*., 2021) for 3, 6, and 9 weeks compared with 0 weeks (chow diet). Mice on NAFLD diet demonstrate weight gain starting at 6 weeks (**Fig S1A**) and induction of ad lib blood glucose levels, indicative of insulin resistance (**Fig S1B**). Gene expression analyses of livers demonstrate elevated levels of the lipogenesis genes *sterol responsive element binding protein-1c (srebf1)*, demonstrating activation of lipid synthesis (**Fig S1C**). In addition, we observe increases in the inflammatory genes *F4/80 (emr1), tumor necrosis factor-α (tnf-α), interleukin-1β (il1β), transforming growth factor-β 1 (tgfβ1)*, and the fibrosis genes *collagen type1 α 1 (col1α1)* and *matrix metalloproteinase 9* (*mmp9)*, suggestive of activation of inflammation and fibrosis transcriptional programs in diet-induced NAFLD (**Fig S1C**). Liver dysfunction was evidenced by a significantly higher liver mass at 9 weeks (**Fig S1D**) and an increase in plasma ALT levels to 35 U/L at 6 weeks, and 60 U/L at 9 weeks (**Fig S1E**). Histological analyses of livers confirmed an increase in lipid accumulation by Oil red O staining (**Fig 1A**) as well as macroscopically larger livers (**Fig 1B**). Next, we isolated parenchymal hepatocytes and non-parenchymal cell populations in the liver using density-gradients (Jung, Zhao and Svensson, 2020). Surprisingly, hepatocytes isolated from mice with established NAFLD demonstrate a large degree of heterogeneity in their ability to accumulate lipids (**Fig 1C**). The cell isolation method was further confirmed by the expression of the hepatocyte markers *pon1, albumin and srebf1* (**Fig 1D**), the highly enriched immune cell markers *il-6, il-1b*, and *emr1* in the non-parenchymal cell fraction (**Fig 1E**), as well as the fibroblast markers *tgfβ1, mmp9, col1α1* and *α-sma* (**Fig 1F**).

**Figure 1.**
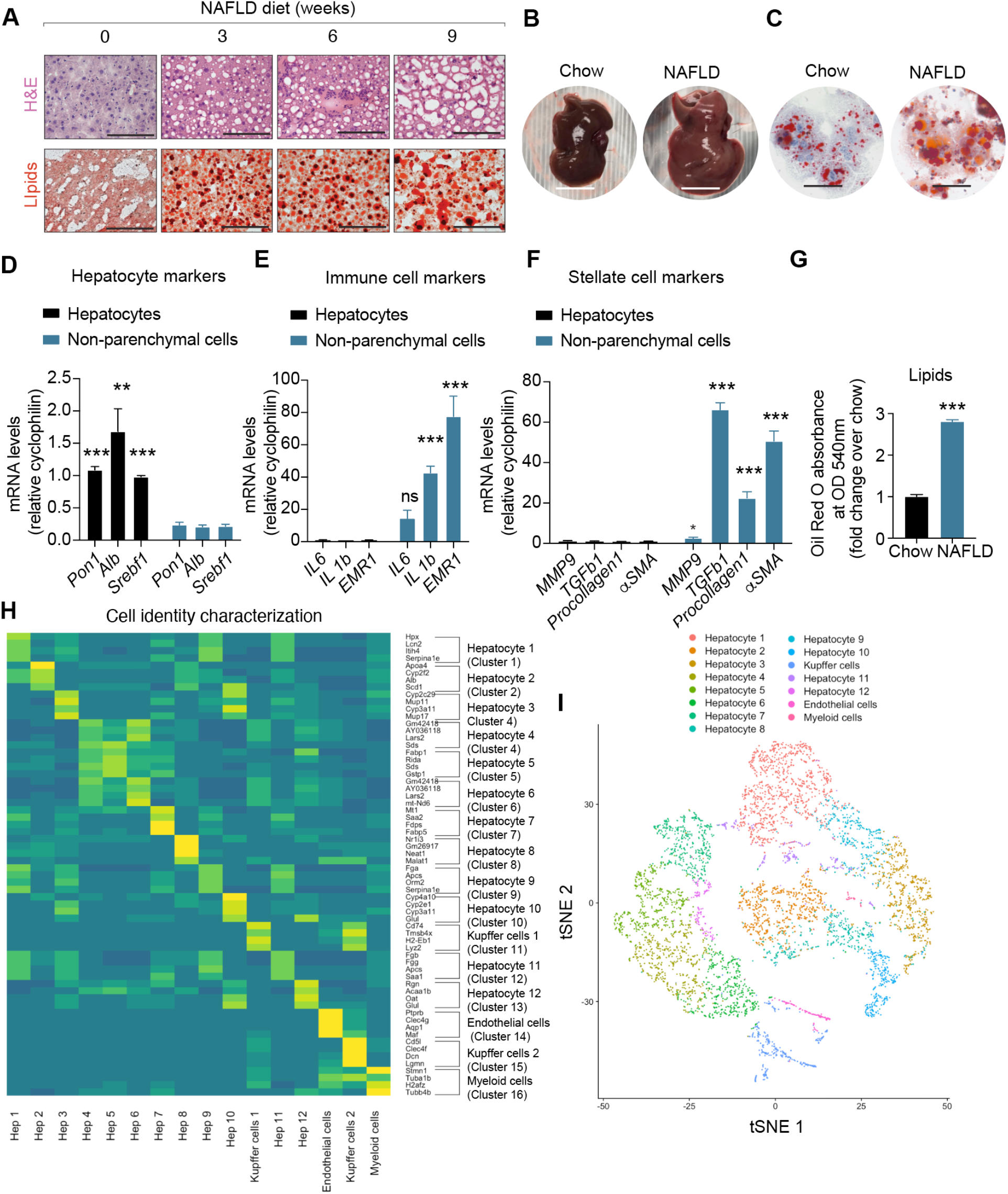
Hepatic steatosis is characterized by high hepatocyte heterogeneity. **A**. Histological analyses using H&E and Oil red O (lipids) in livers from mice fed NAFLD diet for 0, 3, 6 and 9 weeks, *N* = 5 mice per group. Scale bars = 100µm. **B**. Macroscopic photographs of livers from mice fed chow diet or NAFLD diet for 6 weeks. One representative image is shown from an experiment with *N* = 5 mice per group. **C**. Oil red O staining demonstrating cellular accumulation of lipids in hepatocytes isolated from mice fed chow or NAFLD diet for 6 weeks. One representative image is shown from an experiment with *N* = 5 mice per group. **D**. Gene expression analysis of hepatocyte markers in isolated hepatocytes and non-parenchymal cells (*N* = 3 samples/group). (*N* = 3 samples/group, ± S.E.M * p < 0.05, ** p < 0.01, *** p < 0.001 by two-tailed Student’s t-test). **E**. Gene expression analysis of immune cell markers in isolated hepatocytes and non-parenchymal cells (*N* = 3 samples/group). (*N* = 3 samples/group, ± S.E.M * p < 0.05, ** p < 0.01, *** p < 0.001 by two-tailed Student’s t-test). **F**. Gene expression analysis of fibrosis/stellate cell markers in isolated hepatocytes and non-parenchymal cells (*N* = 3 samples/group, ± S.E.M * p < 0.05, ** p < 0.01, *** p < 0.001 by two-tailed Student’s t-test). **G**. Oil red O staining quantification of hepatocytes isolated from chow of NAFLD livers (*N* = 3 samples/group). Data are presented as mean ± S.E.M of biologically independent samples. * p < 0.05, ** p < 0.01, *** p < 0.001 by two-tailed Student’s t-test. **H**. Gene set enrichment criteria for the identification of cell types in the dataset of merged chow and NAFLD clusters identified by Seurat. Values are presented as [log2] fold change. Enrichment was calculated for cells in the cluster over all cells outside the cluster. **I**. Aggregated t-SNE plot of merged chow and NAFLD cell clusters identified by sc-RNA sequencing.

To characterize the molecular changes and heterogeneity at the single-cell level, we next performed single-cell RNA sequencing (scRNA-seq) at 6 weeks, at which time point a >2.5-fold increase in intracellular lipid accumulation was observed (**Fig 1G)**. Freshly isolated hepatocytes and non-parenchymal cells were isolated from chow-or NAFLD-fed mice (*N* = 4 mice/group), combined, and processed for single-cell RNA sequencing library synthesis preparation after confirming cell viability. Importantly, all samples were sequenced in the same round to avoid the need for batch correction. A total of 5932 cells expressing 17,606 genes were included in the analysis following filtering to remove very high (>0.5) mitochondrial genome transcript ratio, genes detected (UMI count >0) in less than three cells, and cells with very small library size (<1000) (see **Methods**). Cells were clustered and visualized using aggregated single-cell expression relationship profiles with t-distributed stochastic neighbor embedding (t-SNE) plots using Seurat (Butler *et al*., 2018a) (**Fig 1H-I**). 16 populations were clustered: 12 distinct hepatocyte populations, and 4 non-parenchymal cell types (**Fig 1H-I**). We identify five main cellular identities in both conditions – hepatocytes, stellate cells, kupffer cells, myeloid and endothelial cells in line with previously described cell types in the liver (**Fig S1F**) (Halpern *et al*., 2017; Ben-Moshe *et al*., 2019). These data confirm that the liver is characterized by high hepatocyte heterogeneity.

### Hepatocytes display dynamic changes during NAFLD progression

To perform more in-depth analyses of the gene signatures in hepatocytes during NAFLD progression, we next clustered cells based on significant cell identity and expression. While kupffer cells, endothelial cells and myeloid cells did not markedly change their expression at the six-week time point (**Fig S2A-C** and **Fig S3**), hepatocytes from NAFLD mice demonstrated large changes in expression as illustrated in the t-SNE plots comparing chow and NAFLD livers (**Fig 2A**). Specifically, we observe distinct gene signatures in the hepatocyte clusters, illustrated in ridgeline plots showing the heterogenous expression of *Glul* (glutamine synthetase) (**Fig 2B**), a gene encoding for a metabolic enzyme abundantly expressed in hepatocytes (Pettinelli *et al*., 2018; Ma *et al*., 2020). Other hepatocyte genes such as *lipocalin-2* (*Lcn2*), *Hpx, Apcs, Gstp1* and *cyp4a10* demonstrate similar heterogeneity in expression across hepatocyte clusters, suggesting distinct cellular states (**Fig 2B**). Furthermore, fibrinogen alpha (*Fga*) is enriched in the chow hepatocyte clusters (**Fig 2C**), while *2-iminibutanoate/2-iminopropanoate deaminase (rida*), and *serine dehydratase (sds)*, two enzymes involved in amino acid metabolism, are enriched in NAFLD hepatocyte clusters (**Fig 2D**). Interestingly, two novel genes, *Gm42418* and *AY036118* were enriched in NAFLD hepatocyte clusters, indicating a potential function during NAFLD (**Fig 2D**). These analyses demonstrate that NAFLD is associated with dramatic changes in lipogenic gene expression that are heterogeneously distributed even within a given cell type.

**Figure 2.**
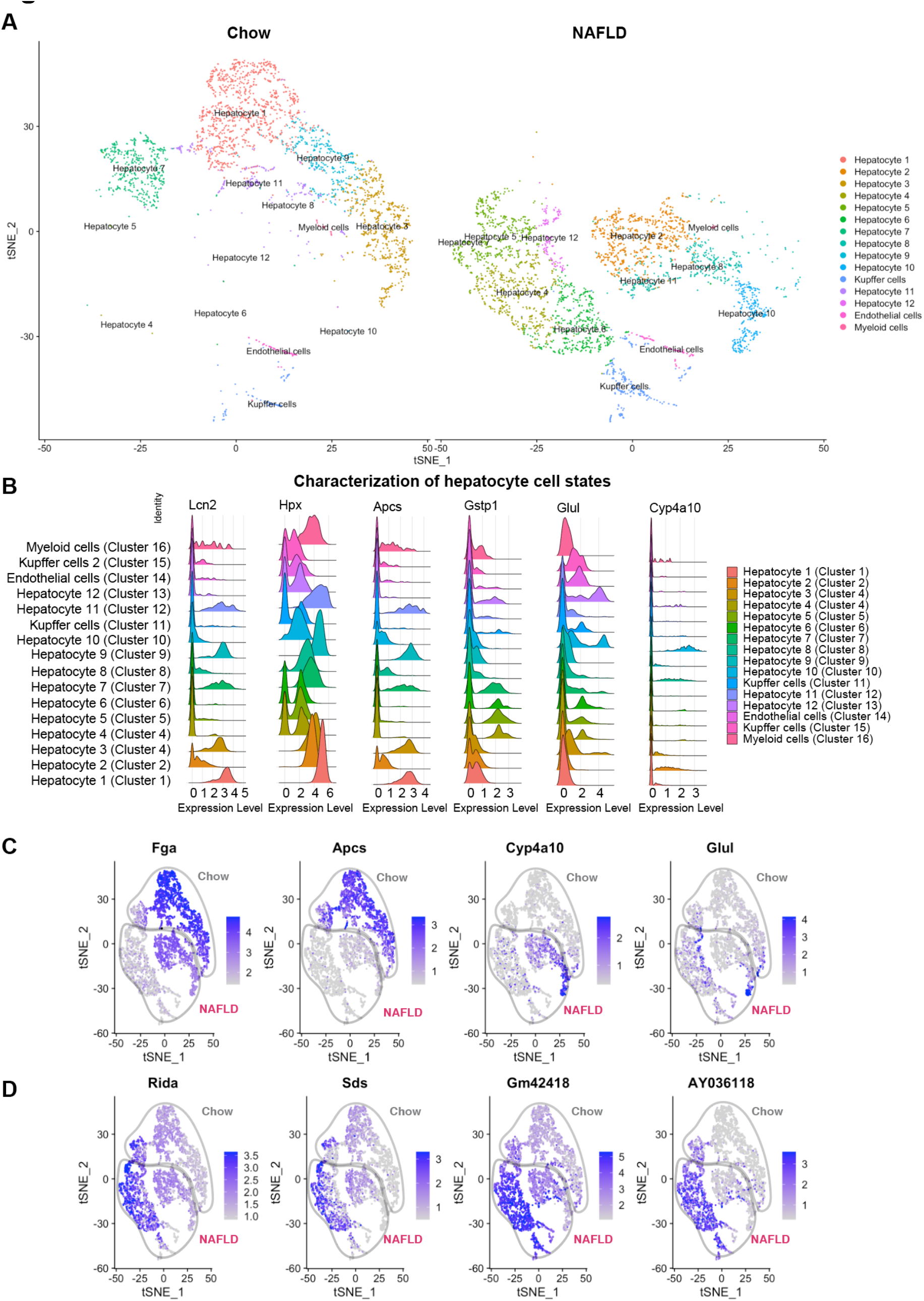
Hepatocytes display dynamic changes during NAFLD progression. **A**. t-SNE plot of cell clusters separated by chow and NAFLD identified by Seurat. **B**. Ridgeline plots of the hepatocyte genes *Lcn2, Hpx, Apcs, Gstp1, Glul*, and *Cyp4a10* across all cell clusters identified by Seurat. Values are presented as log2 fold change. **C**. t-SNE plots of hepatocyte-enriched genes *Fga, Apcs, Cyp4a10*, and *Glul* across chow (grey) and NAFLD (red) groups identified by Seurat. Expression in tSNE plots is shown as normalized transcript counts on a color-coded log2 scale. **D**. t-SNE plots of hepatocyte-enriched genes *Rida, Sds, Gm42418*, and *Ay036118* across chow (grey) and NAFLD (red) groups identified by Seurat. Expression in tSNE plots is shown as normalized transcript counts on a color-coded log2 scale.

### Hepatocyte populations display large heterogeneity in *Srebf1* expression

*Srebf1*, which is under the direct control of both insulin and fructose is one of the most well-described transcriptional drivers of lipogenesis (Huang *et al*., 2010; Softic, Cohen and Kahn, 2016; Gosis *et al*., 2022). Interestingly, under both chow and NAFLD conditions, there were two main clusters defined by high or low expression of *srebp1c* (*srebf1*) (**Fig 3A-B**). When performing single cluster analysis of the *srebp1* expression, two main populations appear to be dominant: *srebp1*^*high*^ (clusters 1, 2, 3, 8, 9, 10) and *srebp1*^*low*^ (clusters 4, 5, 6, 7, 11, 12) (**Fig 3A**). Notably, the high expression of *lxra (nr1h3*) and *chREBP (mlxipl)* in *srebp1*^*high*^ clusters suggests a broad activation of the classical lipid synthesis pathways (**Fig 3A-B**). The general hepatocyte marker albumin (*alb)* was equally expressed across hepatocyte clusters, suggesting that the NAFLD state specifically induces the expression of genes involved in lipid synthesis. This finding was unexpected and led us to question whether these changes in expression could explain the differences in lipid accumulation seen in these hepatocytes. Nuclear srebp1 staining revealed a highly heterogenous expression in both chow and NAFLD when *srebp1*^*high*^ and *srebp1*^*low*^ cells were quantified (**Fig S4A-C**). To correlate lipid accumulation with srebp1 expression in *srebp1*^*high*^ and *srebp1*^*low*^ cells in intact liver sections, we performed co-immunostainings of srebp1 and the lipid droplet dye LipidSpot. Expectedly, only negligible levels of lipid droplets could be detected under chow conditions, while lipid droplet accumulation and increased lipid droplet size were observed in NAFLD at 6 weeks. Notably, nuclear srebp1 protein expression did not co-localize with the areas that demonstrated a high density of lipid droplets (**Fig S4D-E**). These data support that both *srebp1*^*high*^ and *srebp1*^*low*^ cells with robust lipid droplet accumulation in NAFLD are present, suggesting that alternative lipogenic drivers might exist.

**Figure 3.**
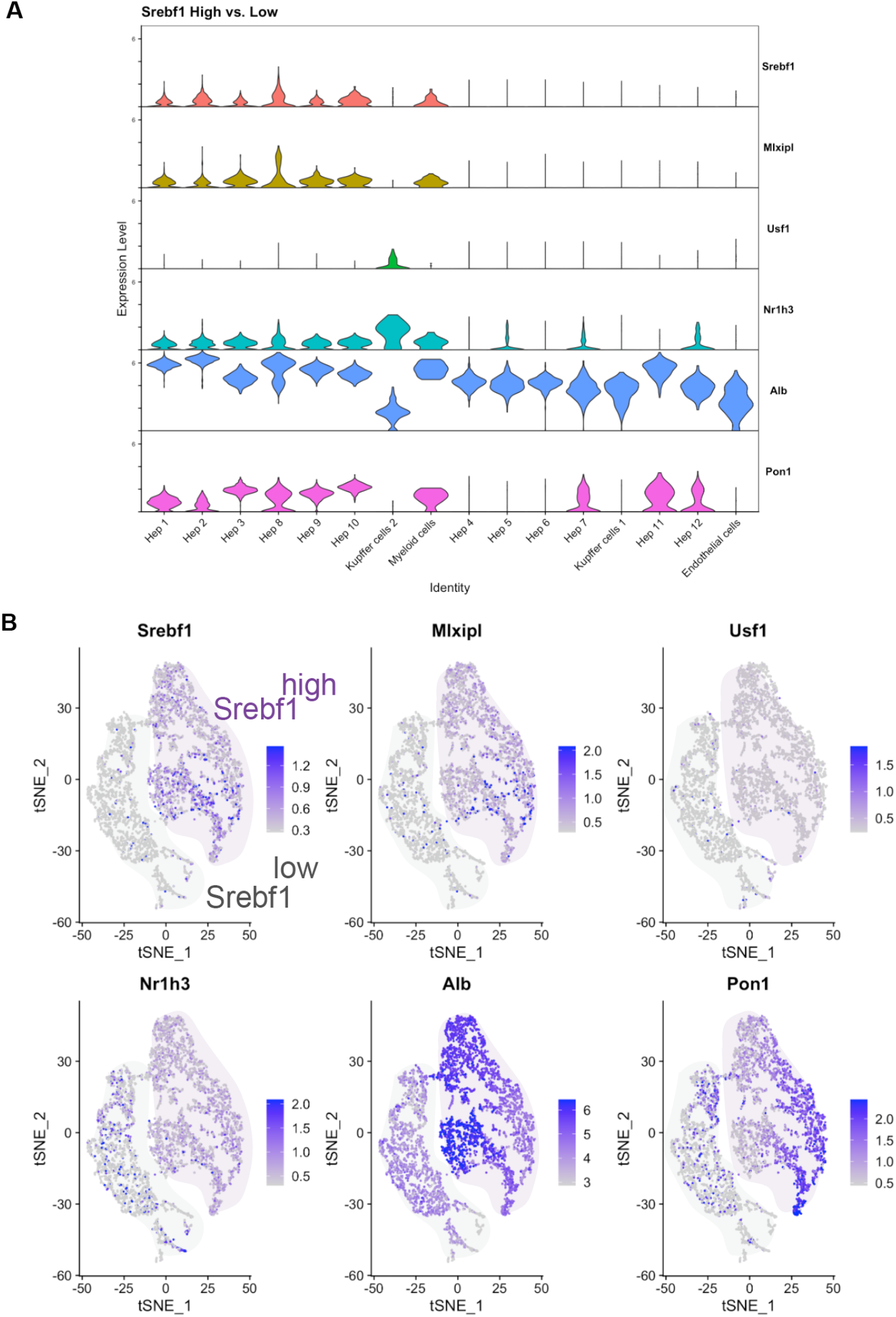
Hepatocyte populations display large heterogeneity in *Srebf1* expression. **A**. Violin plots of *srebf1, chrebp (mlxipl), usf1, lxrα (nr1h3), alb*, and *pon1* in cell clusters. Values are log2 expression levels. **B**. t-SNE plots of srebf1^high^ and srebf1^low^ clusters displaying the cell type localization of *srebf1, chrebp (mlxipl), usf1, lxrα (nr1h3), alb*, and *pon1*.

### Identification of NR1I3 (CAR) as a NAFLD-enriched nuclear receptor in mice

To identify other transcriptional regulators that might play a role in NAFLD, we next compared the liver transcriptome between human and mouse livers using a previously published human dataset from Bader *et. al* in combination with the current mouse single-RNA sequencing dataset. Out of 21,128 detected genes in either dataset, 12,398 genes were present in both humans and mice (**Fig S5**) and correlated well in expression (**Fig 4A-B**). In the mouse dataset, we next analyzed the differentially expressed genes between NAFLD and chow across all hepatocyte populations (**Fig 4C and Table S1**) and found that among the top NAFLD-enriched genes, Nr1i3 (nuclear receptor subfamily 1, group I, member 3), or constitutive androstane receptor (CAR) was the only nuclear receptor (**Fig 4D and Table S1**). These data are consistent with prior studies of the global CAR-KO mice, which are protected from hyperlipidemia and diet-induced fibrosis (Yamazaki *et al*., 2007; Maglich, Lobe and Moore, 2009). However, the CAR agonist TCPOBOP attenuates steatohepatitis in mice fed a methionine-choline deficient (MCD) diet (Dong *et al*., 2009). In addition, the role of CAR in human NAFLD and steatohepatitis is unclear.

**Figure 4.**
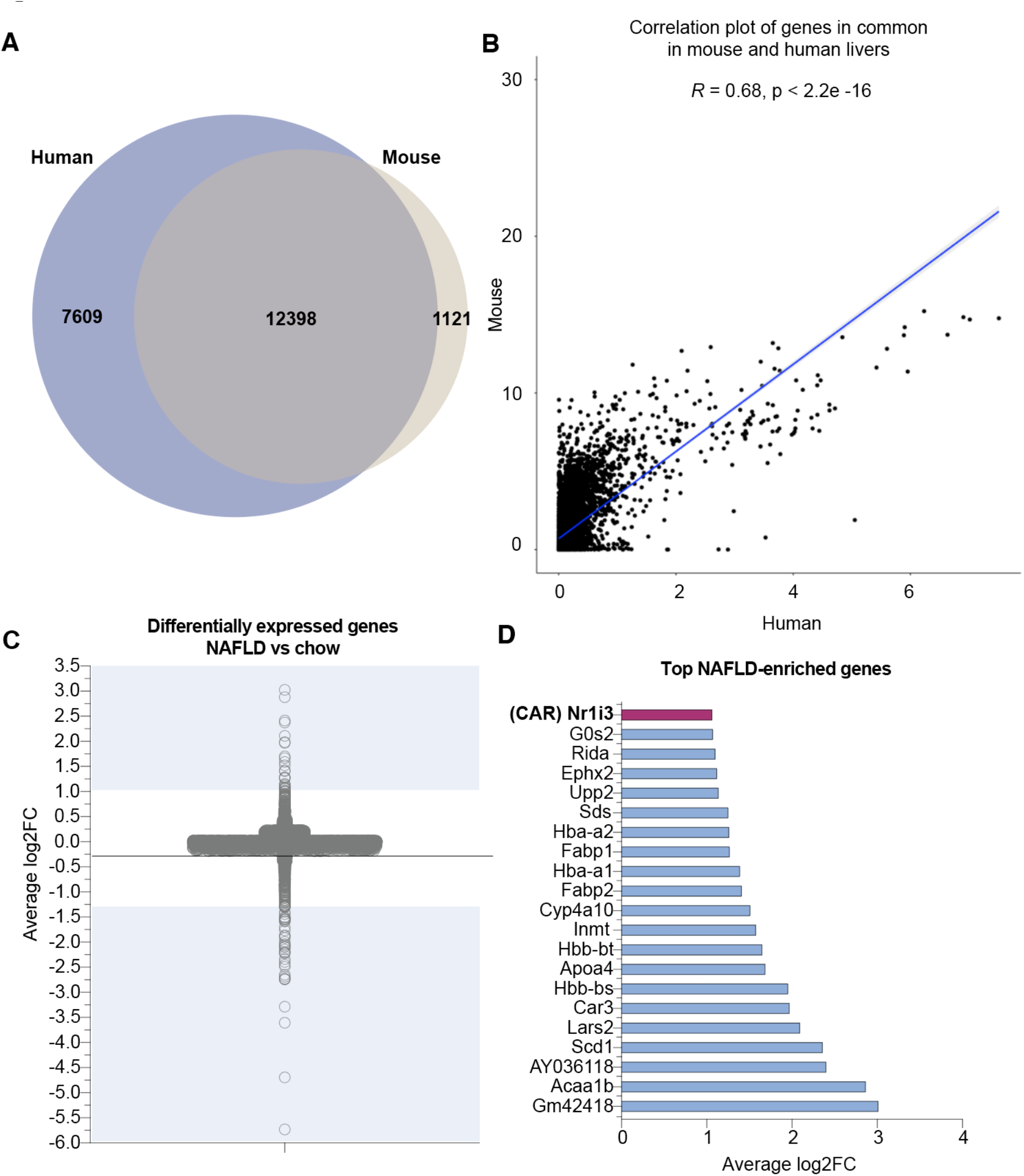
Identification of NR1I3 (CAR) as a NAFLD-enriched nuclear receptor in mice. **A**. Overlap of human and mouse genes detected across two independent single-cell RNA seq datasets. **B**. Correlation plot of genes in common in mouse and human livers (Pearson correlation, *R* 0.68, p < 2.2 × 10^−16^). **C**. Differentially expressed genes in NAFLD vs chow aggregated across all clusters identified in the single-cell RNA sequencing dataset expressed as average [log2 fold change (FC)]. **D**. Top genes enriched in NAFLD vs chow from (C) expressed as average [log2FC].

### A predictive model independently identifies CAR as a human NASH gene

Independently, using Artificial Intelligence (AI) embedding methods, we created a predictive model to identify novel NAFLD/NASH-related genes and pathways with the hypothesis that the predicted genes would modulate relevant liver biology. To identify novel disease associated genes, we built a predictive NASH-model based on the proximity of the genes to functional modules in our embedded map of protein-protein interactions (PPIs) (see **Methods**). We performed 10-fold cross validation and systematically varied the parameters of the model. We tested 308 NASH genes from the BeFree dataset from DisGeNET (Piñero *et al*., 2020), and 134 NASH genes identified experimentally (**Table S2**). Our model achieves AUC of 0.82 and 0.83, respectively in recovering these NASH disease genes (**Fig S6**). In seeking novel key genes, we were particularly interested in transcription factors that might drive the steatohepatitis processes of lipogenesis, fibrosis, and others. Using the above NASH-model, CAR (human gene name NR1I3) was independently identified as a transcription factor with high likelihood to associate with NASH in humans (**Fig 5**). From a group of 1200 transcription factors, 22 are computationally identified as liver expressed, and 12 out of 22 are associated with NASH. Interestingly, CAR interacts with four critical functional modules: Cholesterol Homeostasis, Bile Acid Metabolism, Fatty Acid Metabolism, and Estrogen Response (**Fig 5**).

**Figure 5.**
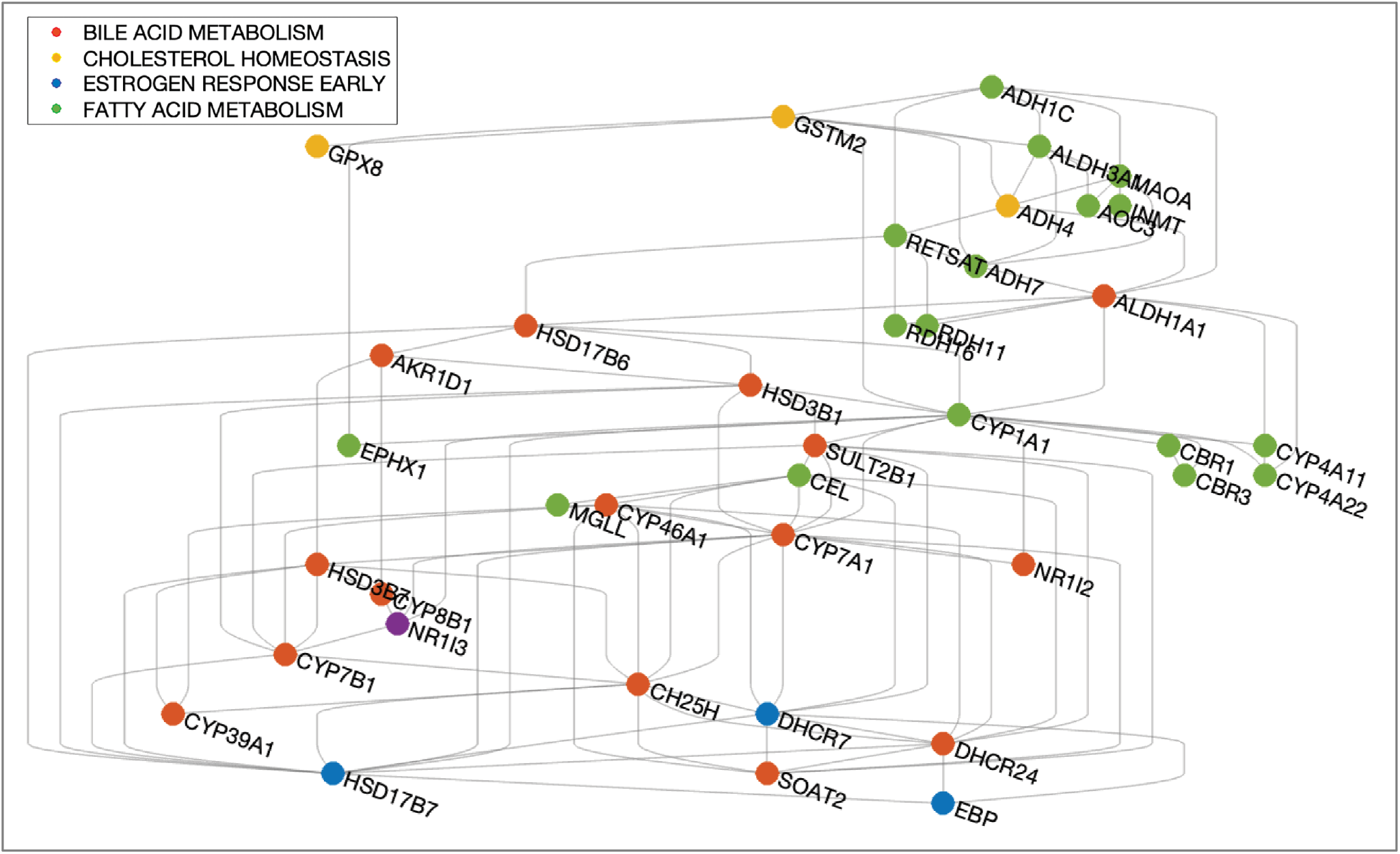
A predictive model independently identifies CAR as a human NASH gene. Direct and indirect connections between NR1I3 and 51 genes. A predictive NASH-model independently identifies NR1I3 as a human NASH gene (purple dot). It has strong computationally predicted associations with 51 genes in four critical NASH functional modules (colored as indicated in the legend).

### Nuclear CAR localization correlates with steatohepatitis in humans

We next sought to determine whether CAR expression levels correlated with NAFLD in human liver samples. 26 liver sections from 13 patients with no histopathological abnormality and 13 patients with histological steatohepatitis were included in the analysis (see **Methods**). There was no significant difference in body mass index (BMI) (**Fig 6A**), age (**Fig 6B**), or sex (**Fig 6C**) between the groups. The steatohepatitis group had higher triglyceride levels (**Fig 6D**), alanine aminotransferase activity (ALT) (**Fig 6E**), and aspartate transaminase (AST) activity (**Fig 6F**) and were diagnosed with histologic grades of 1-2 (**Fig 6G**) and histologic stage of 0-3 (**Fig 6H**). Using a specific validated antibody for CAR (Petryszak *et al*., 2016), we quantified the pixel intensity (**Fig 6I**) of CAR nuclear staining across the 26 liver samples. We found that the CAR protein expression levels positively correlated with steatohepatitis (**Fig 6J**), while the gene expression in a separate cohort did not significantly correlate (**Fig S7A-H**). Representative microscopic images validate the lipid droplet size and nuclear localization of CAR (**Fig 6K**), with no staining in the negative control (**Fig S7I**). Lastly, correlation analyses demonstrate that ALT and AST significantly correlated with CAR protein expression (**Fig 6L-M**), while BMI did not significantly correlate with CAR levels (**Fig 6N**). These results show that nuclear CAR expression correlates with steatohepatitis in humans.

**Figure 6.**
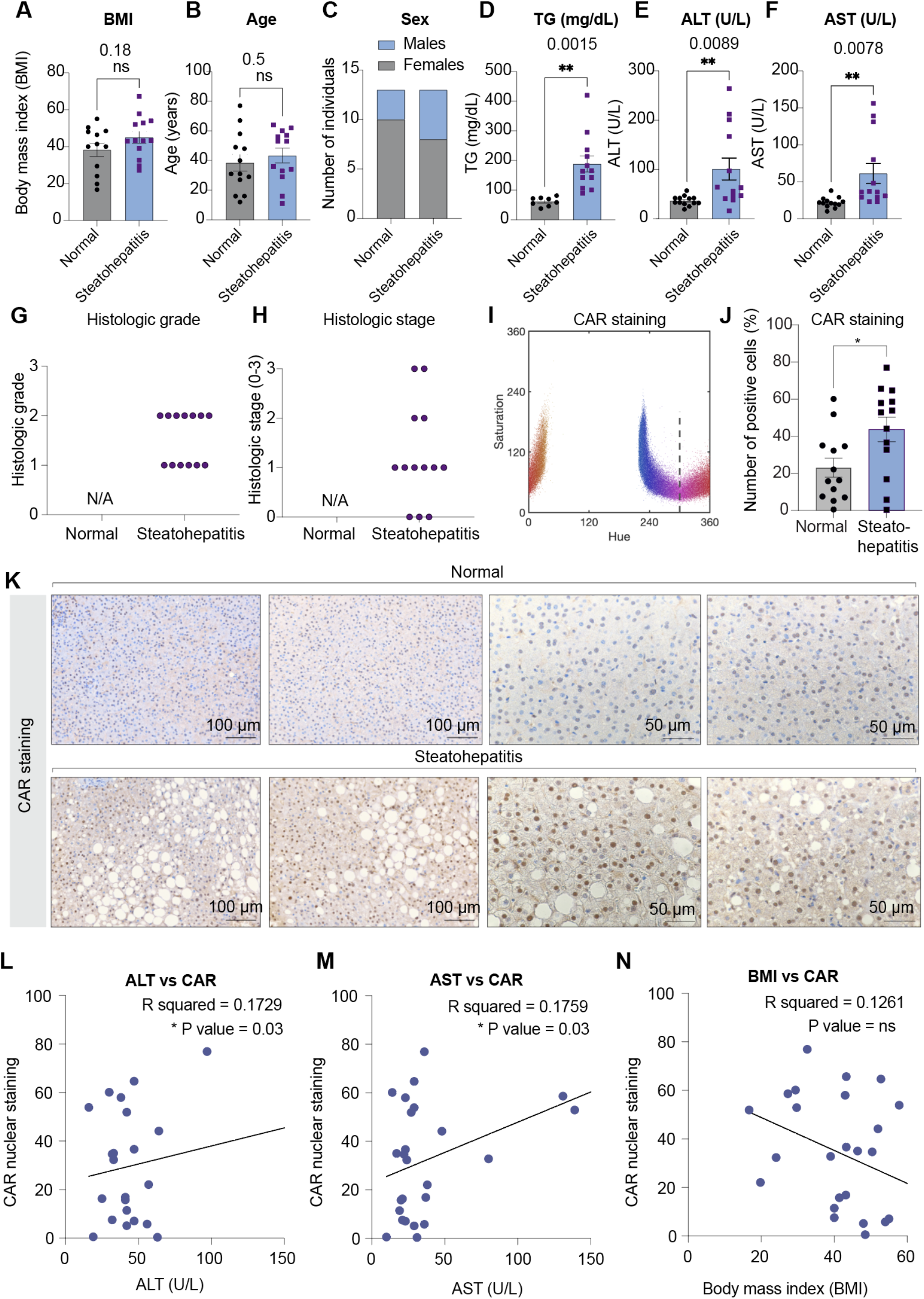
Nuclear CAR localization correlates with steatohepatitis in humans. **A**. Body mass index (BMI) of individuals with normal livers vs. livers with steatohepatitis (*N* = 13 patient samples per group, S.E.M. *P < 0.05 by two-tailed Student’s t-test). **B**. Age of individuals with normal livers vs. livers with steatohepatitis (*N* = 13 patient samples per group, S.E.M. *P < 0.05 by two-tailed Student’s t-test). **C**. Sex of individuals with normal livers vs. livers with steatohepatitis (*N* = 13 patient samples per group, S.E.M. *P < 0.05 by two-tailed Student’s t-test). **D**. Triglyceride (TG) levels in mg/dL of individuals with normal livers vs. livers with steatohepatitis (*N* = 13 patient samples per group, S.E.M. *P < 0.05 by two-tailed Student’s t-test). **E**. Alanine transferase (ALT) activity in U/L of individuals with normal livers vs. livers with steatohepatitis (*N* = 13 patient samples per group, S.E.M. *P < 0.05 by two-tailed Student’s t-test). **F**. Aspartate transaminase AST) activity (in U/L of individuals with normal livers vs. livers with steatohepatitis (*N* = 13 patient samples per group, S.E.M. *P < 0.05 by two-tailed Student’s t-test). **G**. Histological grade of individuals with normal livers vs. livers with steatohepatitis (*N* = 13 patient samples per group, S.E.M. *P < 0.05 by two-tailed Student’s t-test). **H**. Histological stage of individuals with normal livers vs. livers with steatohepatitis (*N* = 13 patient samples per group, S.E.M. *P < 0.05 by two-tailed Student’s t-test). **I**. Segmentation of positive (Red) and negative (blue) cells based on pixel intensity in the nuclei from all images combined to determine the positive threshold for quantification. Each dot represents one cell. **J**. Quantification of the % number of CAR-positive nuclei of individuals with normal livers vs. livers with steatohepatitis (*N* = 13 patient samples per condition, 4 images per sample). Data are presented as S.E.M. *P < 0.05 by two-tailed Student’s t-test. **K**. Representative histological images of CAR staining (*N* = 13 patient samples per condition, 4 images per sample) in normal livers or livers with steatohepatitis. Bar scale : 100 µm or 50 µm. **L**. Correlation plot of CAR levels and ALT levels (U/L) across 26 individuals. *P value < 0.05. **M**. Correlation plot of CAR levels and AST levels (U/L) across 26 individuals. *P value < 0.05 **N**. Correlation plot of CAR levels and BMI across 26 individuals. *P*-value = ns (non-significant).

## DISCUSSION

Our study uncovers several aspects of liver biology that were previously unknown. First, we find that hepatocytes are highly heterogeneous in their capacity to store lipids and metabolic profiles. By capturing these lipid-laden hepatocytes for single-cell analyses, we find that this heterogeneity is associated with lipid metabolism in a subset of hepatocytes. Elegant previous work analyzing transcriptional changes in hepatocytes isolated from mice fed a 60 % high-fat diet showed that hepatic steatosis sensitizes cells to hepatocyte inflammation (Sheng, Jiang and Rui, 2013), suggesting that the stored lipid content is a principal determinant of hepatocyte function. How the lipid heterogeneity of hepatocytes contributes to NAFLD progression will be an area for future exploration.

Second, we find that high *srebp1* expression is not directly associated with higher lipid accumulation, indicating the involvement of other driving factors in a subset of hepatocytes. The finding of a subset of genes in hepatocytes co-expressed with *srebf1* warrants further investigations into the mechanisms regulating lipid metabolism *in vivo*.

Third, using experimental and AI predictive models, we identify CAR as a gene highly associated with steatohepatitis in mice and humans. CAR is abundantly expressed in hepatocytes, but the reported role of CAR in non-alcoholic fatty liver disease is controversial (di Masi *et al*., 2009; Dong *et al*., 2009; Maglich, Lobe and Moore, 2009; Lynch *et al*., 2014). We find here that CAR is localized to the nucleus and overexpressed in patient livers diagnosed with steatohepatitis. Future studies should assess the association and functional correlation between CAR and NAFLD/NASH, as well as explore targeting CAR using specific agonists. Moving forward, it will be important to investigate whether CAR regulates lipogenesis, inflammation, or fibrosis during NASH using relevant models.

Collectively, our results uncover an unexpected heterogeneity in hepatocyte steatosis and identify novel lipogenic cell populations. In conclusion, this study facilitates the discovery of previously unrelated genes involved in lipid metabolism, which may be used to better understand how to target fatty liver disease.

## Limitations of the study

There are limitations to this work. While our study demonstrates heterogenous hepatocyte cell state signature in NAFLD, the functions of these distinct hepatocyte populations remain to be determined. In addition, single-cell analyses of hepatocytes should be complemented with single nuclei analyses to confirm the hepatocyte cell states identified in this paper.

## Author contributions

Conceptualization, L.C., Y.J. K.J.S.; Methodology, L.C., Y.J. S.B.N., K.J.S.; Validation, Y.J., M.Z., L.C., K.J.S.; Formal Analysis, L.C., Y.J., S.B.N., K.J.S; Investigation, L.C., Y.J., M.Z., T.L., L.C., S.P.; Resources, G.C., R.A., A, K-K, H. Y-J., K.J.S.; Writing – Original Draft, L.C., K.J.S.; Writing – Review & Editing, L.C., M.Z., T.L., R.A., K.J.S.; Funding Acquisition, K.J.S.; Supervision, K.J.S.

## Acknowledgements

K.J.S. was supported by NIH grants DK125260, DK111916, the Stanford Diabetes Research Center P30DK116074, the Jacob Churg Foundation, the McCormick and Gabilan Award, the Weintz Family COVID-19 research fund, American Heart Association (AHA), the Stanford School of Medicine, and the Stanford Cardiovascular Institute (CVI). M.Z. was supported by the American Heart Association (AHA) postdoctoral fellowship (905674). L.C. was supported by Stanford School of Medicine Dean’s Postdoctoral Fellowship. S.B.N. was supported by the Novo Nordisk Foundation (grant award NNF20OC0059462) and the Stanford Bio-X Program. reNEW was also supported by the Novo Nordisk Foundation (grant award NNF21CC0073729). This work used the Genome Sequencing Service Center by Stanford Center for Genomics and Personalized Medicine Sequencing Center, supported by the grant award NIH S10OD020141, and the Diabetes Genomics and Analysis Core of the Stanford Diabetes Research Center supported by grant award NIH/NIDDK P30DK116074. We thank Pratima Nallagatla for assistance with bioinformatic analyses and Aila Karioja-Kallio for the collection of human liver samples. We thank the Stanford University Pathology Histology core for histological processing and the Pathology Department for microscopy equipment.

## STAR Methods

### RESOURCE AVAILABILITY

#### Lead Contact

Further information and requests for resources and reagents should be directed to and will be fulfilled by the Lead Contact, Dr. Katrin J. Svensson.

#### Materials availability

All reagents used in this study are commercially available.

#### Data and code availability

All raw and processed single-cell RNA sequencing data have been deposited to Gene Expression Omnibus (GSE210501). The script for immunohistochemistry quantification is available at https://github.com/Svensson-Lab/Coassolo2022. All information required to reanalyze the data is reported in this paper.

### EXPERIMENTAL MODEL AND SUBJECT DETAILS

#### Human samples

The human liver samples used for histological characterization of CAR levels were obtained at Stanford University under IRB protocol #58373 using excess/archival material. The subjects were identified by searching the pathology archive database for liver disease diagnoses. Exclusion criteria were liver cancers and tumors as identified by information stored in the pathology database. Clinical and laboratory data were obtained by retrieving existing data in the electronic medical record. The human liver cDNA samples were obtained from NAFLD/non-NAFLD patients from Hannele Yki-Jarvinen, Finland. The human single-cell RNA sequencing data was re-analyzed from a previously published dataset by Macparland *et al*. (MacParland *et al*., 2018).

#### Animals

Animal experiments were performed per procedures approved by the Institutional Animal Care and Use Committee of the Stanford Animal Care and Use Committee (APLAC) protocol #32982. C57BL/6J mice were purchased from the Jackson Laboratory (#000664). Unless otherwise stated, mice were housed in a temperature-controlled (20-22°C) room on a 12-hour light/dark cycle. All experiments were performed with age-matched male mice housed in groups of five unless stated otherwise. For single-cell RNA sequencing experiments, 12-week-old male mice were fed with either chow diet (Envigo, #2018) or NAFLD diet (40 % fat, 20 % kcal fructose, and 2 % cholesterol, #D09100310 ResearchDiets) for 6 weeks and sacrificed at 18 weeks of age.

#### Isolation of cells for single-cell RNA sequencing

All cells were cultured in a humidified atmosphere containing 5% CO2 at 37°C. Primary hepatocyte isolation was performed as previously described (Jung, Zhao and Svensson, 2020). 12-week-old male C57BL/6J mice fed with either chow diet or NAFLD diet for 6 weeks were sacrificed. The inferior vena cava was cannulated with a 25G needle connected to tubing for perfusion with 20 ml of 37 °C pre-heated pH 7.4 HBSS buffer (#14175-095, Gibco) containing 5.4mM KCl, 30mM Sodium bicarbonate and 0.285 mM EDTA at a rate of 1 ml per minute. After 2 minutes of perfusion, the hepatic portal vein was cut. After perfusion, 20 ml of liver digestion medium containing 1mg/ml collagenase type IV ((#C5138, Sigma), 10% FBS and 1mM HEPES was added to the perfusion buffer tube. After 7-10 minutes, the liver was dissected and transferred to a petri dish and mechanically dissociated by gently swirling the tissue in 10 ml Williams E medium containing Glutamax (#112-033-101, Quality Biological), 10% FBS, 2mM sodium pyruvate, 1uM dexamethasone and 0.1uM insulin. The dissociated liver cells were filtered through a 70 µm cell strainer and centrifuged at 50 x*g* for 4 minutes to separate hepatocytes and non-parenchymal cells (NPC). Supernatants (NPCs) were transferred to a fresh 50 ml tube and the pellet (hepatocytes) was washed by adding 25ml of Williams E medium and centrifuged at 20 x*g* for 3 minutes to reduce the number of erythrocytes. and livers were perfused in HBSS buffer (#14175-095, Gibco) supplemented with 0.4 g/L KCl, 1 g/L glucose, 2.1 g/L sodium bicarbonate, and 0.2 g/L EDTA for 3 minutes, followed by Collagenase (#C5138, Sigma) digestion at 37 ?. Cells were dissociated from the digested livers and hepatocytes were suspended in Williams Medium E (#112-033-101, Quality Biological) supplemented with 10% FBS, 2 mM sodium pyruvate, 1 μM dexamethasone, and 100 nM insulin (plating medium). The cell suspension was filtered through a 70 µm strainer and centrifuged at 50 x*g* for 3 minutes. For isolation of hepatocytes, cell pellets were resuspended in plating medium and mixed with 90 % or 25 % Percoll (#P1644, Sigma) followed by centrifugation at 100 x*g* for 10 minutes and 50 x*g* for 3 minutes. For isolation of NPCs, cell suspensions were centrifuged at 100 x*g* for 5 minutes to reduce the number of erythrocytes. NPCs were isolated and washed as follows: 300 x*g* for 7 minutes, 650 x*g* for 4 minutes, 240 x*g* for 5 minutes, and 650 x*g* for 4 minutes. Pellets were then combined with hepatocytes. Cells were combined as one sample (chow and NAFLD, *N* = 4 mice/group) and processed for single-cell RNA sequencing library synthesis preparation after confirming cell numbers and viability. For culture of hepatocytes and non-parenchymal cells, cell pellets were resuspended in plating medium. 4 hours after seeding on collagen-coated plates, cells were washed with PBS, followed by the addition of maintenance medium Williams E supplemented with 0.2% BSA, 2 mM sodium pyruvate, 0.1μM dexamethasone.

#### Library preparation, single-cell RNA sequencing and data preprocessing

All samples were sequenced in the same round to avoid the need for batch correction. The libraries were prepared using the 10X Genomics 3’ version 3 single cell gene expression kit. Then they were sequenced on the Illumina HiSeq 4000 with a 2×101 bp reads. Dual indexed libraries of isolated mouse liver cells were then pooled and sequenced on an Illumina HiSeq 4000 sequencer at the Stanford Genome Sequencing Service Center in a 100-bp paired-end configuration. The read structure was dual indexed sequencing run with Read 1 starting from a Read 1 being 28 bases including cell barcode and unique molecular identifier (UMI), index i7 of 10 bases, index i5 of 10 bases and Read 2 being 90 bases containing transcript information. The libraries were processed and decomplexed using the pipeline from 10x Genomics Cell Ranger 3.1.0 (Weisenfeld *et al*., 2017).

#### Cluster identification and expression analysis of single-cell RNA sequencing data

The data was analyzed using Seurat (Butler *et al*., 2018a) in R Studio. Raw count tables were loaded into Seurat for both datasets (chow and NAFLD) and the Seurat “merge” function was applied to perform pooled analysis. After removing cells that were either less than 200 genes or more than 50,000 genes and filtering out over 5% mitochondrial content, gene expression was normalized by global-scaling normalization method, “LogNormalize”, merged and clustered following the standard Seurat package procedures (Butler *et al*., 2018b; Stuart *et al*., 2019). The combined dataset of chow and NAFLD mice liver identified populations consistent with prior reports (Suo *et al*., 2018; Franzén, Gan and Björkegren, 2019; Zhang *et al*., 2019). The cluster definition heatmap illustrates three representative markers of gene expressions shown in log2 value for each cluster to define the population. For comparisons between mouse and human sc-RNA sequencing data, the Rshiny app provided by the Bader lab was used to generate and analyze the hepatocyte clusters. In Bader *et al*, clusters 1, 3, 4, 6, 14, and 15 were identified as hepatocytes. The following analyses were performed: (1) scatter plot for all mouse and human genes (12,509 genes found in common between 17,606 genes in the mouse and 20,007 genes in the human datasets) expressed as average gene expression across all samples in the human dataset (8444 cells) vs. mouse (5932 cells, NAFLD and chow), (2) venn diagram of overlap in mouse and human genes, and (3) extraction of the clustering information for the human dataset to determine if genes were heterogeneously expressed in the human hepatocyte clusters. P values were adjusted using Bonferroni correction for multiple testing.

#### AI embedding models for predicting pathogenesis genes

To identify novel disease associated genes, we have built predictive models based on the proximity of the genes to functional modules in our embedded map of PPIs as described previously (Liu *et al*., 2022). We applied node2vec to capture network topology features of the 14,707 genes (from STRING-2019) in a 64-dimensional embedding space (Szklarczyk *et al*., 2019). Node2vec is a graph algorithm that uses the local connectivity around a gene to summarize its interactions in a low dimensional space (Grover and Leskovec, 2016); neighboring nodes have similar interactions and are “close” in the embedding space, as measured by cosine distance. We then estimated a gene’s role in pathogenesis by computing its embedding distance from a set of 220 functional modules (47 immune response modules from ImmProt (Rieckmann *et al*., 2017), 50 signaling pathway modules from MsigDB (Liberzon *et al*., 2015), 123 metabolic modules from Human Metabolic Reaction Database (Wishart *et al*., 2007) to those of known disease genes. Given a module consisting of a set of genes, we summed the individual gene embedding vectors to generate a summary vector for the functional module. Cosine similarity measures proximity between gene embeddings and module vectors. For a given gene, we can calculate its similarity to each of the above functional modules; the vector of values describes the genes functional relationships. Given a gold standard set of genes involved in disease, we can predict additional disease genes using machine learning on these embedded representations of functional relationships. As a proof of concept, we collected 70 NASH genes from DisGeNET (Piñero *et al*., 2020) as a gold-standard list. We identified the functional modules that are closest to these 70 NASH genes. We used a Linear SVM to create a classifier for defining additional genes with a similar profile as the known 70. A held out set of randomly chosen genes were used as negative examples. We performed 10-fold cross validation and systematically varied the parameters of the model.

#### Immunohistochemistry

Immunohistochemistry on mouse livers was performed on OCT-embedded 6 μm thick frozen sections. Tissue sections were fixed in 3% formalin in PBS for 1h at 20°C. For hematoxylin and eosin (H&E) staining, slides were stained with hematoxylin, washed with water and 95% ethanol, and stained with eosin for 30 min. Sections were then dehydrated with ethanol and xylene and mounted with mounting medium. Trichrome staining was performed according to the manufacturer’s instructions using Trichrome Stain Kit (Sigma, #HT15). For CAR quantification, immunostaining was performed on paraffin-embedded human liver tissues. In brief, the paraffin blocks were sliced into 5-μM thick sections, deparaffinized with xylene and rehydrated with decreasing concentrations of ethanol in water. Antigen retrieval was achieved by incubating slides in sodium citrate buffer (pH 6.0) for 25 min at 95°C, followed by 20 min of cooling at room temperature. Endogenous peroxidases were quenched by incubating the slides in 3% hydrogen peroxide for 10 min. The sections were then washed with phosphate-buffered saline (PBS) for 5 min. Endogenous avidin and biotin were blocked using a blocking kit according to manufacturer instructions (Vector Laboratories, Burlington, ON, Canada). Primary antibody (1/100, #CF805306, ThermoFisher) was applied for 2h at room temperature in a humidified chamber. After rinsing the slides in PBS, they were incubated in polymer-HRP secondary antibody (Vector Laboratories) for 30 min at room temperature. After washing with PBS for 5 minutes, the slides were incubated with Vectastain ABC reagent (Vector Laboratories) for 30 min. After washing with PBS for five minutes, color development was achieved by applying diaminobenzidine tetrahydrochloride (DAB) solution (Vector Laboratories) for two minutes. After washing in distilled water, the sections were counterstained with hematoxylin and blue in ammonia water, dehydrated through ethanol and xylene, and cover-slipped using a xylene-based mounting medium. All slides were reviewed at the same time with a Nikon eclipse e1000 upright light microscope and images were captured with a Nikon Digital Sight DS-Fi1 color camera and Spot Advanced software.

#### CAR quantification

Segmentation was done independently for each image by converting an RGB image into grayscale and stretching the converted pixel intensities such that the top 1 % and the bottom 1 % of the pixels were saturated. A threshold was introduced at 40 % of the maximum pixel intensity. Any pixel with a grayscale intensity below that threshold was considered part of a cell nucleus. To improve the segmentation further, a flood-filled operation and a 5-pixel disk-shaped morphological opening were applied. The average RGB pixel color inside each segmented nuclei was calculated, which was considered the most likely color of that cell nuclei. To classify nuclei into positive and negative cells, the RGB pixel color was converted into HSV space. Any cell nuclei with a hue inside the interval [120; 300] was considered negative; otherwise, the cell was considered positive. The script is available at https://github.com/Svensson-Lab/Coassolo2022.

#### Oil red O staining and quantification

For Oil red O staining on slides or in cells, samples were fixed with 3 % formalin in PBS, washed twice with water, incubated with 60 % isopropanol for 5 min and then incubated with premixed Oil Red O solution (Sigma, #O1395) at the ratio of 3:2 of Oil Red O:H2O for 20 min followed by washing with H2O. For quantification of Oil Red O staining in cells, Oil Red O staining was extracted by adding 0.5 ml isopropanol to stained cells in a 24-well plate and transferred to a 96-well clear flat-bottomed plate. Absorbance was read at 540 nm. Values are expressed as absorbance normalized by cell number.

#### Immunofluorescence staining and quantification

For Srebp1 (Santa Cruz, sc-13551) and LipidSpot dye (Biotium, 70069, 1:1000 dilution in PBS) co-staining, the staining was performed sequentially. The Srebp1 antibody (1:200 dilution in PBS/1 % BSA) was applied for 24h at 4 °C, followed by a secondary fluorescent antibody (Thermo Fisher Scientific, A11001, 1:500 dilution in PBS/1 % BSA) for 1h. LipidSpot dye was then applied for 2h on the same liver tissue, according to the manufacturer’s protocol. The fluorescent images of Srebp1 and LipidSpot were taken using a Leica SP8 confocal microscope using a 63x objective. Quantification of nuclear-localized Srebp1 was performed on immunofluorescence images using the ImageJ software. Briefly, 5 representative pictures (90µm x 90 µm) of lipid positive and negative areas were selected from the 6 weeks of NAFLD diet mice. The median intensity of Srebp1 staining was measured in the Srebp1^high^ and Srebp1^low^ cells. Srebp1 was interpreted as positive based on the set threshold. Appropriate positive and negative controls were reviewed at the same time. The co-localized area (µm^2^) of Srebp1 and DAPI staining was quantified. The values are expressed as percentage of cells with co-localized nuclei in DAPI-positive cells. The statistical analysis was performed by comparisons between lipid positive and negative areas using two tailed Student’s t-test. P-values represent ****: p <0.001, ***: p <0.01, **: p <0.05, *: p <0.1, non-significant: p>0.1.

#### Plasma Alanine Transaminase (ALT) activity measurement

Animals fed with either chow or NAFLD diet were sacrificed, and blood was drawn by cardiac puncture and collected in heparinized tubes. Blood was centrifuged at 6000 x*g* for 10 minutes at 4 °C to collect plasma and the supernatant was transferred to a new tube and kept at -80 °C. ALT activity measurements and analyses were performed according to the manufacturer’s protocol using Alanine Transaminase Colorimetric Activity Assay Kit (Cayman, #700260). Briefly, 150 µl of substrates, 20 µl of cofactor and 20 µl of 2x diluted plasma samples were loaded onto 96 well plates in duplicates including positive controls that were provided in the kit. The plates were incubated for 15 minutes at 37 °C. After incubation, the reactions were initiated by adding 20 µl of ALT initiator followed by immediate measurement of the absorbance at 340 nm every 10 minutes at 37 °C. The absolute difference in absorbance value between two time points of was divided by the time point difference, which then was multiplied by the extinction coefficient of 0.21/(4.11 × 0.02) and the dilution factor. Values are expressed as units (U)/L.

#### RNA expression analysis

Total RNA from cultured cells or tissues was isolated using TRIzol (#15596018 Thermo Fisher Scientific) and Rneasy mini kits (# 74104 Qiagen). RNA was reverse transcribed using the ABI high-capacity cDNA synthesis kit. For qRT-PCR analysis, cDNA, primers and SYBR-green fluorescent dye (ABI) were used. Relative mRNA expression was determined by normalization with *cyclophilin* levels using the ΔΔCt method. The primer sequences used are listed in the Key Resource Table.

#### Statistical analysis of cellular and animal experiments

All values in graphs are presented as mean +/- S.E.M. P-values represent ****: p <0.001, ***: p <0.01, **: p <0.05, *: p <0.1, non-significant: p>0.1. Two-way ANOVA was used for repeated measures (* p < 0.05, ** p <0.01, *** p < 0.001). Student’s t test was used for single comparisons. Values for *N* represent biological replicates for cultured cell experiments or individual animals for *in vivo* experiments. Specific details for *N* values are noted in each figure legend. For cellular assays, *N* corresponds to the number of experimental replicates using cells isolated from individual mice. Each cellular experiment using primary cells was repeated using at least two cohorts of mice. For animal experiments, *N* corresponds to the number of animals per condition. Mice were randomly assigned to treatment groups for *in vivo* studies. Each animal experiment was repeated using at least two cohorts of mice.

## SUPPLEMENTARY INFORMATION

**Figure S1.**
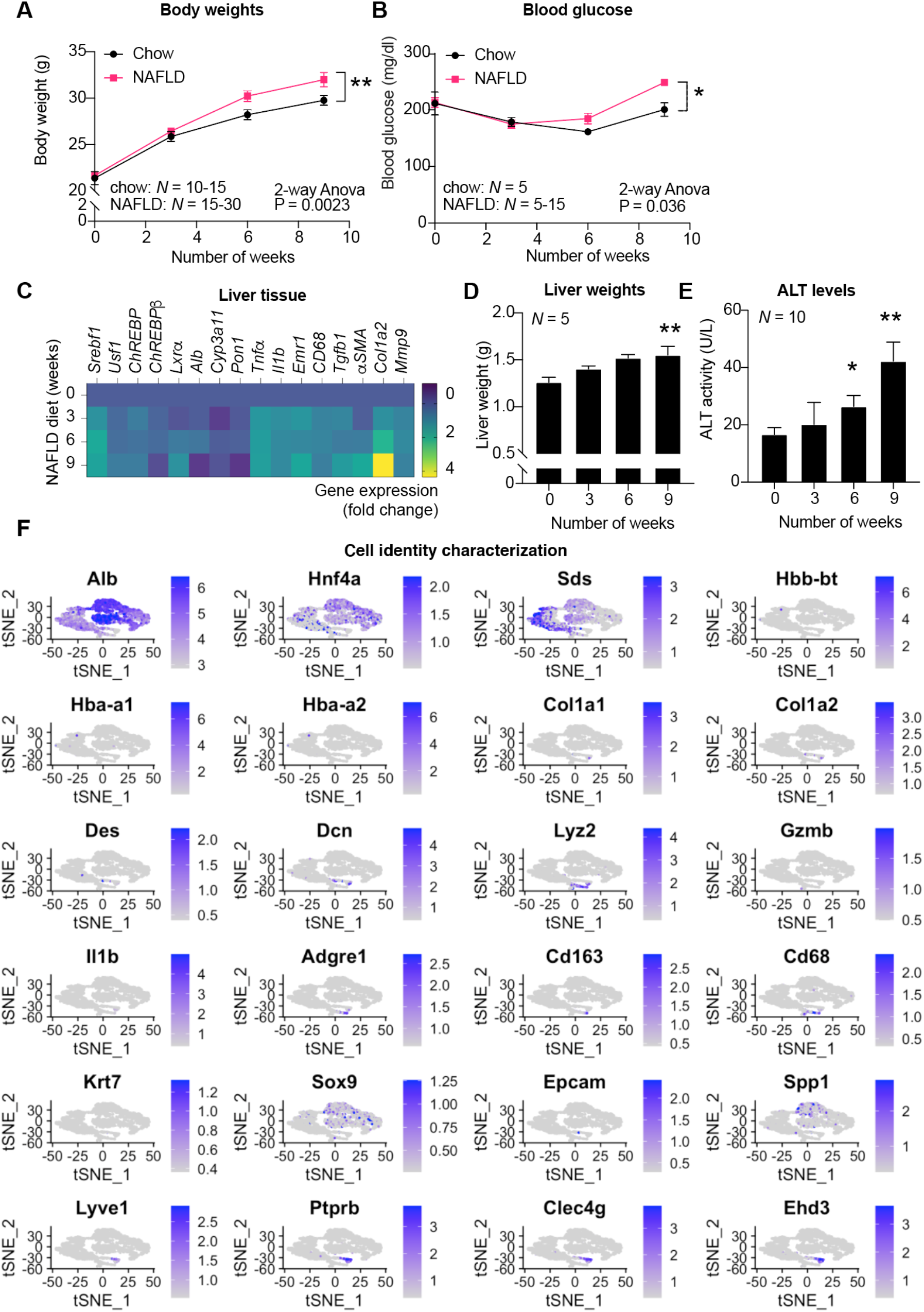
Characterization of the NAFLD mouse model and single-cell analysis. **A**. Body weights of mice fed NAFLD diet for 0, 3, 6 and 9 weeks combined from two independent cohorts (chow, 0 weeks *N* = 10; chow 3 weeks, *N* = 10; chow 6 weeks *N* = 15, chow 9 weeks, *N* =15; NAFLD 0 weeks, *N* = 30; NAFLD 3 weeks, *N* = 30; NAFLD 6 weeks, *N* = 30; NAFLD 9 weeks, *N* = 15). Data are presented as mean ± S.E.M of biologically independent samples. * p < 0.05, ** p < 0.01, *** p < 0.001 by two-way Anova. **B**. Blood glucose levels in mice fed NAFLD diet for 0, 3, 6 and 9 weeks (chow, *N* = 5, NAFLD 0 weeks *N* = 20; NAFLD 3 weeks, *N* = 20; NAFLD 6 weeks *N* = 10, NAFLD 9 weeks, *N* = 5). Data are presented as mean ± S.E.M of biologically independent samples. * p < 0.05, ** p < 0.01, *** p < 0.001 by two-way Anova. **C**. Gene expression analysis of lipid synthesis, inflammation, and fibrosis genes in livers from mice fed NAFLD diet for 0, 3, 6 and 9 weeks, *N* = 5 mice per group. **D**. Liver weights isolated from mice fed NAFLD diet for 0, 3, 6 and 9 weeks, *N* = 5 mice per group. Data are presented as mean ± S.E.M of biologically independent samples. * p < 0.05, ** p < 0.01, *** p < 0.001 by student’s two-tailed t-test. **E**. Plasma ALT levels in mice fed NAFLD diet for 0, 3, 6 and 9 weeks, *N* = 10 mice per group. Data are presented as mean ± S.E.M of biologically independent samples. * p < 0.05, ** p < 0.01, *** p < 0.001 by student’s two-tailed t-test. **F**. t-SNE plots of cell-type specific markers used to classify cell types in Figure 1H.

**Figure S2.**
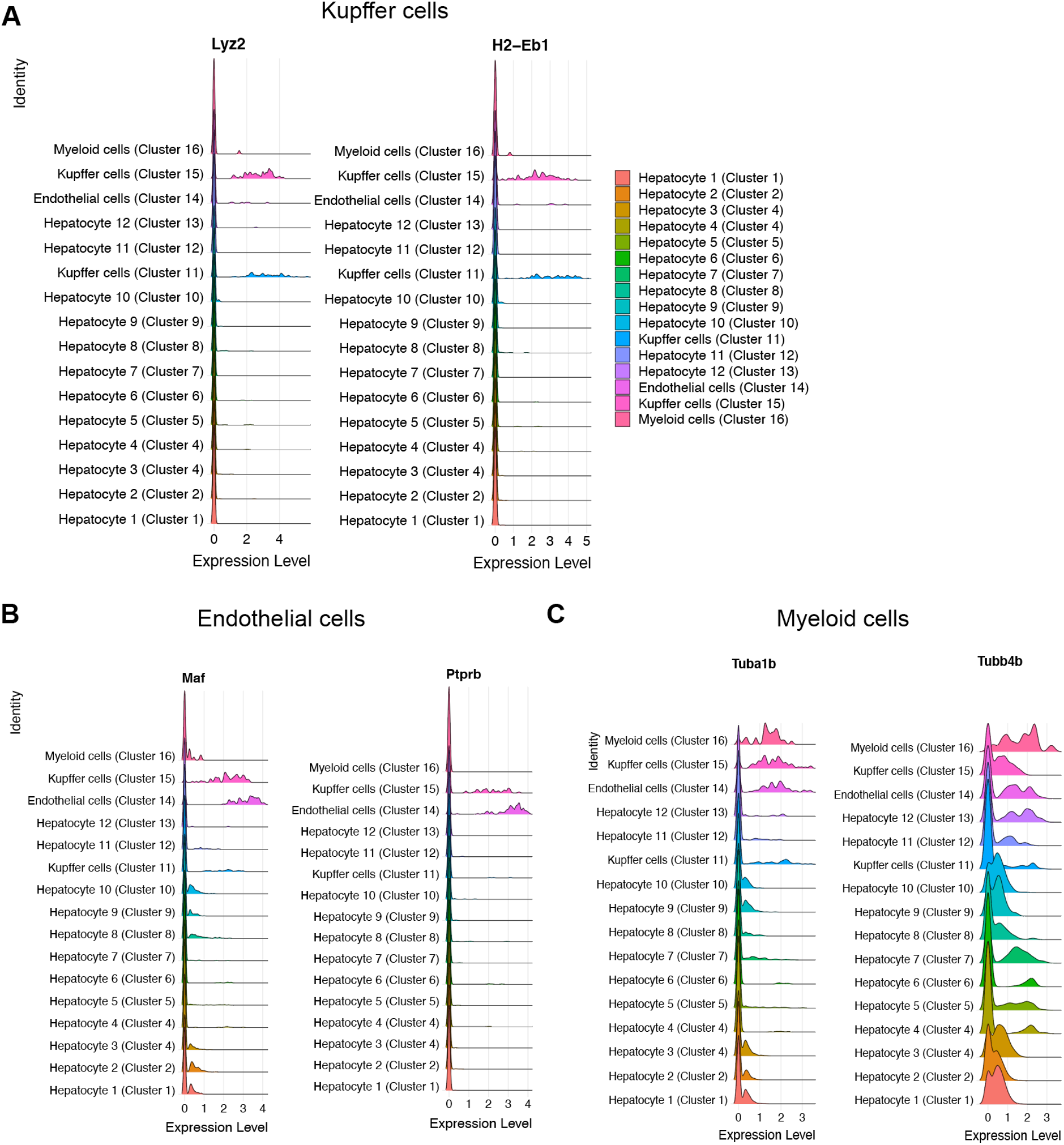
Characterization of other cell types in the liver. **A**. Ridgeline plots of the Kuppfer cell markers *Lyz2* and *H2-Eb1* across all cell clusters identified by Seurat. **B**. Ridgeline plots of the Endothelial cell markers *Maf* and *Ptprb* across all cell clusters identified by Seurat. **C**. Ridgeline plots of the Myeloid cell markers *Tuba1b* and *Tubb4b* across all cell clusters identified by Seurat.

**Figure S3.**
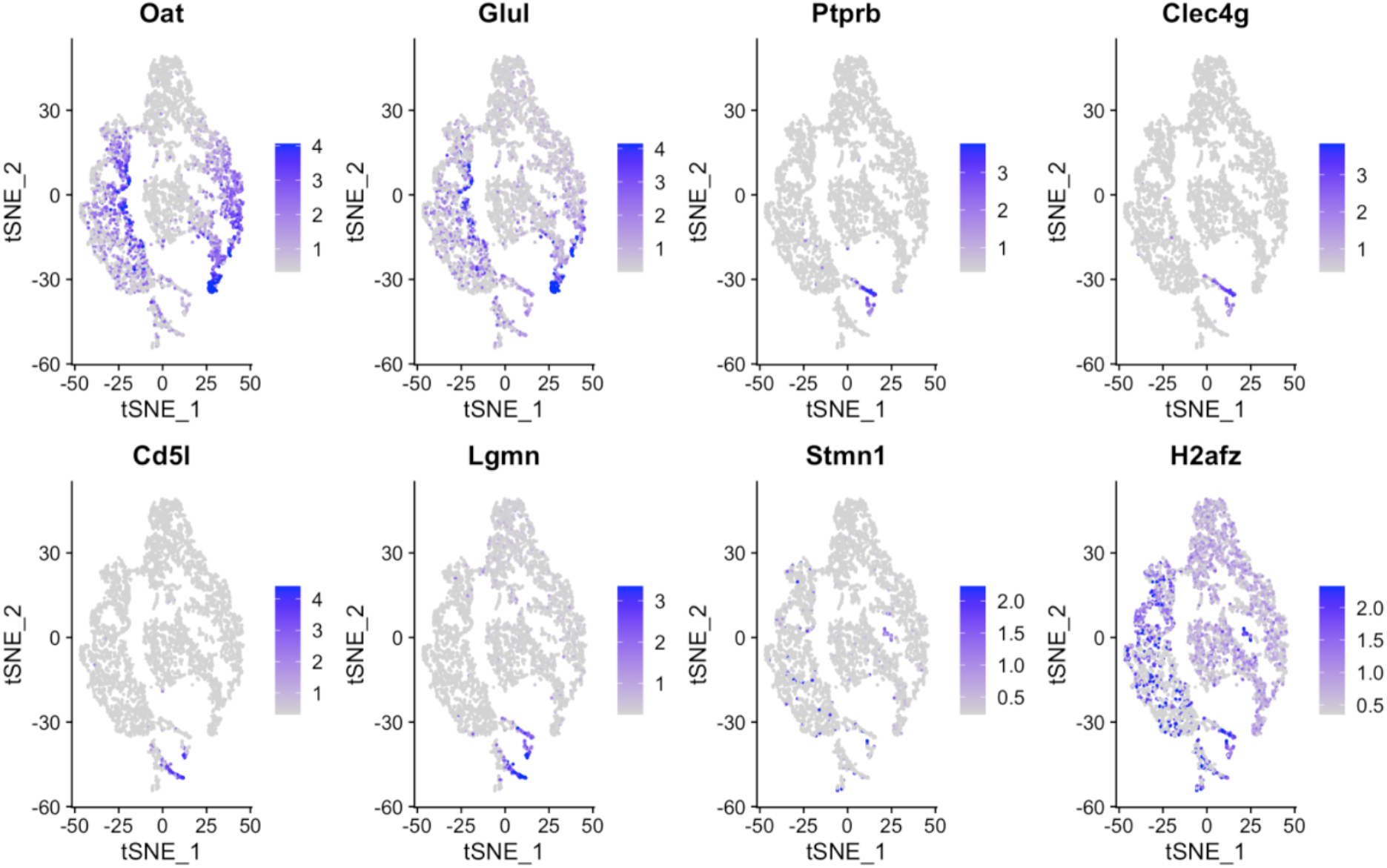
Additional characterization of cell types. t-SNE plots of the additional cell type specific markers for identity characterization in Fig 1H.

**Figure S4.**
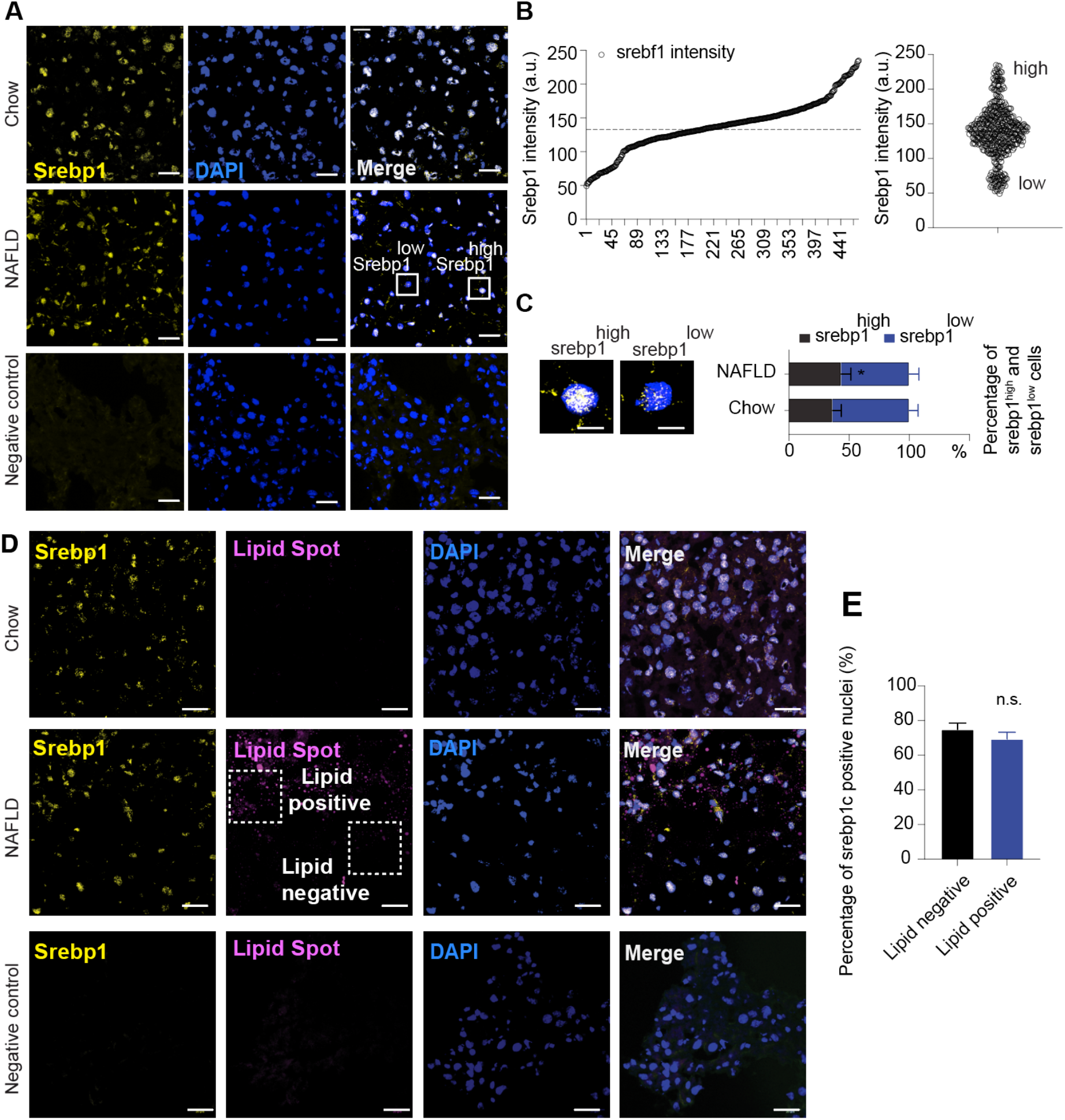
Hepatic Srebp1 levels are heterogeneous and do not correlate with lipid levels. **A**. Confocal microscopy analyses of Srebp1c (yellow), and nuclei (blue) in liver tissues in chow and NAFLD (6 weeks) livers. Scale bar denotes 25 µm. **B**. Threshold for Srebp1c^high^ cells and Srebp1c^low^ cells based on intensity. **C**. Representative images of Srebp1c^high^ and Srebp1c^low^ hepatocytes are shown by enlarged pictures and quantification of Srebp1c^high^ and Srebp1c^low^ populations (N = 10 images per group). **D**. Confocal microscopy analyses of Srebp1c (yellow), lipid droplets (magenta) and nuclei (blue) in liver tissues in chow and NAFLD (6 weeks) livers. Scale bar denotes 25 µm. **E**. Srebp1c positive cells were quantified from five representative images of NAFLD livers. Bar graph shows the fraction of Srebp1c positive nuclei within a lipid negative or lipid positive areas. All data are presented as mean ± S.E.M of 5 images and by two-tailed Student’s t-test.

**Figure S5.**
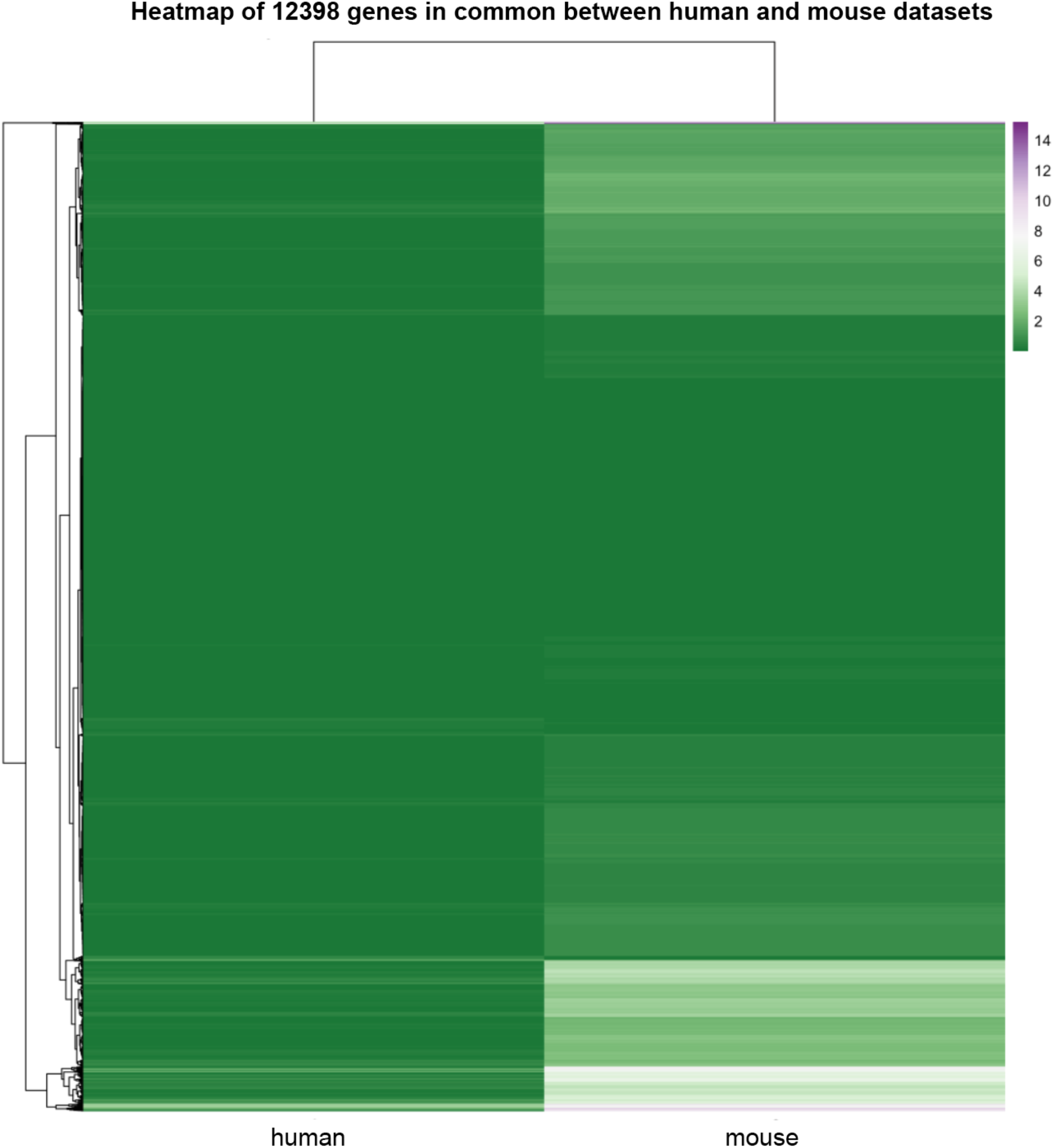
Common hepatic genes between mouse and human datasets. Heatmap comparing genes expressed in mice and humans. Values are expressed as log2 expression.

**Figure S6.**
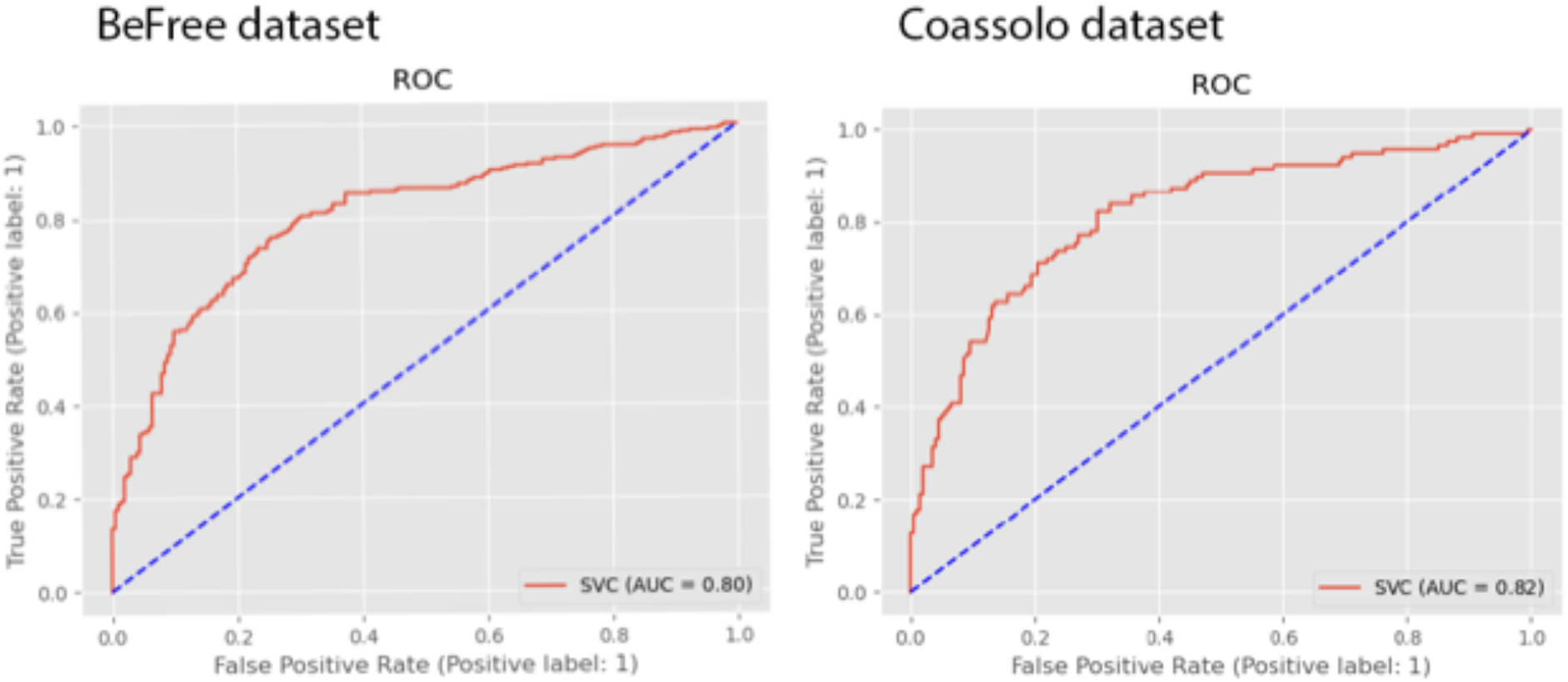
The Area Under the Curve (AUC) performance of the NASH model in predicting NASH-associated genes for two datasets.

**Figure S7.**
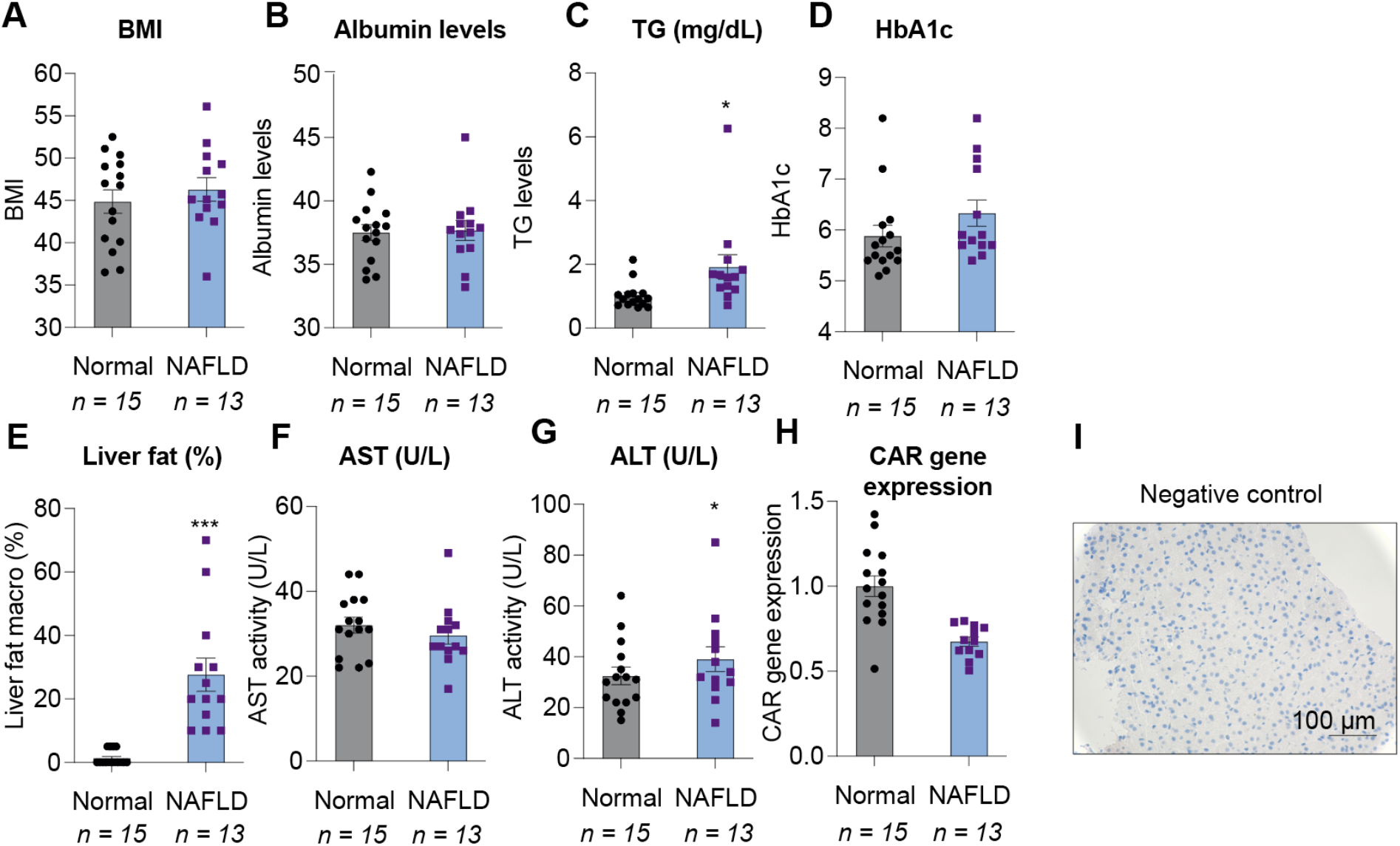
Total CAR mRNA expression does not correlate with NAFLD in humans. **A**. Body mass index (BMI) of individuals with normal livers vs. livers with NAFLD (*N* = 15 normal, *N* = 13 NAFLD samples per group, S.E.M., P value = ns, non-significant). **B**. Albumin levels in individuals with normal livers vs. livers with NAFLD (*N* = 15 normal, *N* = 13 NAFLD samples per group, S.E.M., P value = ns, non-significant). **C**. Triglyceride levels (mg/dL) in individuals with normal livers vs. livers with NAFLD (*N* = 15 normal, *N* = 13 NAFLD samples per group, S.E.M. *P < 0.05 by two-tailed Student’s t-test). **D**. HbA1c levels in individuals with normal livers vs. livers with NAFLD (*N* = 15 normal, *N* = 13 NAFLD samples per group, S.E.M., P value = ns, non-significant). **E**. Liver fat % in individuals with normal livers vs. livers with NAFLD (*N* = 15 normal, *N* = 13 NAFLD samples per group, S.E.M. ***P < 0.001 by two-tailed Student’s t-test). **F**. AST levels (U/L) in individuals with normal livers vs. livers with NAFLD (*N* = 15 normal, *N* = 13 NAFLD samples per group, S.E.M., P value = ns, non-significant). **G**. ALT levels (U/L) in individuals with normal livers vs. livers with NAFLD (*N* = 15 normal, *N* = 13 NAFLD samples per group, S.E.M., P value = ns, non-significant). **H**. Hepatic CAR mRNA expression levels in individuals with normal livers vs. livers with NAFLD (*N* = 15 normal, *N* = 13 NAFLD samples per group, S.E.M., P value = ns, non-significant). **I**. Negative control staining corresponding to CAR staining in Figure 5. Scale bar 100 µm.

